# Distinct Neural Signatures of Outcome Monitoring following Selection and Execution Errors

**DOI:** 10.1101/853317

**Authors:** Faisal Mushtaq, Samuel D. McDougle, Matt P. Craddock, Darius E. Parvin, Jack Brookes, Alexandre Schaefer, Mark Mon-Williams, Jordan A. Taylor, Richard B. Ivry

## Abstract

Losing a point in tennis could result from poor shot selection or faulty stroke execution. To explore how the brain responds to these different types of errors, we examined feedback-locked EEG activity while participants completed a modified version of a standard three-armed bandit probabilistic reward task. Our task framed unrewarded outcomes as either the result of errors of selection or errors of execution. We examined whether amplitude of a medial frontal negativity (the Feedback-Related Negativity; FRN) was sensitive to the different forms of error attribution. Consistent with previous reports, selection errors elicited a large FRN relative to rewards and amplitude of this signal correlated behavioral adjustment following these errors. A different pattern was observed in response to execution errors. These outcomes produced a larger FRN, a frontocentral attenuation in activity preceding this component, and a subsequent enhanced error positivity in parietal sites. Notably, the only correlations with behavioral adjustment were with the early frontocentral attenuation and amplitude of the parietal signal; FRN differences between execution errors and rewarded trials did not correlate with subsequent changes in behavior. Our findings highlight distinct neural correlates of selection and execution error processing, providing insight into how the brain responds to the different classes of error that determine future action.

## Introduction

When an action fails to produce the desired goal, there is a “credit assignment” problem to resolve: Did the lack of reward occur because the wrong course of action was selected, or was it because the selected action was poorly executed? Consider a tennis player who, mid-game, must determine whether losing the last point was the result of selecting the wrong action or executing the action poorly. The player might have attempted a lob rather than the required passing shot, an error in action selection. Alternatively, a lob might have been appropriate but hit with insufficient force, an error in motor execution.

Reinforcement learning presents a framework for understanding adaptive behavior through trial and error interactions with the environment. According to numerous models (e.g. temporal difference learning; Sutton & Barto, 1998), the discrepancy between expected and actual outcomes, the reward prediction error, provides a learning signal that allows an agent to refine its predictions and update its action selection policy. But what happens when a negative prediction error could arise from either poor action selection or poor response execution?

To address this question, McDougle et al. (2016) used a “bandit” task in which participants chose between two stimuli to maximize reward. In one condition, choices were made using a standard button-press method, a situation in which the negative prediction errors on unrewarded trials were attributed to poor action selection (given the negligible demands on motor execution). In a second condition, choices were made by reaching to the desired bandit. Here, unrewarded trials were attributed to movement execution errors. In the latter condition, participants strongly discounted the negative prediction errors on unrewarded trials relative to the former condition. The authors hypothesized that errors credited to the motor execution system block value updating in the action selection system. Consistent with this hypothesis, McDougle et al. (2019) reported that reward prediction error coding in the human striatum was attenuated following execution errors, relative to selection errors. Differences between responses to selection and execution errors have been attributed to a greater sense of “agency” in the latter, with participants’ choice biases indicating a belief that they can reduce execution errors by making more accurate movements (Parvin et al., 2018).

A window into the processes that underlie outcome monitoring is offered through the discovery of the Feedback-Related Negativity (FRN), a negative deflection in the EEG first identified following the presentation of feedback indicating incorrect responses (Miltner et al., 1997). Following its identification, the component quickly became the subject of intense investigation as a marker signaling gains and losses (Gehring & Willoughby, 2002) and outcomes that are worse than expected (Holroyd et al., 2006). The most prominent explanation of its significance, the “reinforcement learning theory of the error-related negativity” (RL-ERN; Holroyd & Coles, 2002) holds that the component (and its response-locked variant, the Error-Related Negativity, the ERN) indexes the activity of signals from the midbrain dopamine that are conveyed to the anterior cingulate cortex for adaptive modification of behaviour (Holroyd & Coles, 2002; Holroyd & Umemoto, 2016). Recent developments reveal that much of the variation in this component is driven by a positive going component (a Reward Positivity; RewP) responding to outcomes that are better than expected (Foti et al., 2011; Holroyd et al., 2008; Proudfit, 2015). Irrespective of whether this signal is framed as a feedback negativity or reward positivity (here, we refer to this component as the FRN-the most widely label), there is a consensus, as indicated by a meta-analysis of 55 datasets (Sambrook & Goslin, 2015), that it is sensitive to reward prediction error.

The FRN’s sensitivity to errors of action is more contentious. A series of experiments (Krigolson et al., 2008; Krigolson & Holroyd, 2006, 2007a) contrasting high level (goal-attainment) errors, variously operationalized as a failure to reach a target (Krigolson et al., 2008; Krigolson & Holroyd, 2007a), avoid a collision (Krigolson & Holroyd, 2006, 2007b), and the erroneous selection of the wrong hand or force (de Bruijn et al., 2003) with low-level errors (i.e. mismatch between actual and intended motor command), concluded that the latter do not elicit a FRN. Instead, reflecting a hierarchical error processing system (Krigolson & Holroyd, 2006), these motor errors are proposed to be mediated within posterior parietal cortex (Desmurget et al., 1999, 2001; Diedrichsen, 2005). Further elaborations indicated that the FRN may only be generated for action errors that cannot be corrected (Krigolson et al., 2008; Krigolson & Holroyd, 2007a), indicating a binary high level coding of outcomes in the FRN (i.e. signaling whether the goal was achieved or not). In line with this, a recent experiment isolating reward-based and sensory error-based motor adaptation reported a FRN in response to binary reward feedback, but not sensory error feedback-which instead generated a P300 (Palidis et al., 2019). Previous work on the P300’s sensitivity to “low level” motor execution errors led to the proposal that this later parietally distributed component might reflect the revision of an internal forward model in posterior parietal cortex (Krigolson & Holroyd, 2007a).

A contrasting set of results suggest that the FRN (and its response-locked variant, the ERN) may in fact be sensitive to motor errors and reflect more than binary coding of outcomes, with evidence showing that it scales with the magnitude of error during sensorimotor adaptation (Anguera et al., 2009) and correlates with the size of hand-path deviations following externally perturbation to target reaches (Torrecillos et al., 2014). These findings are more in line with a growing body of work suggesting that the FRN indexes a general salience prediction error (Oliveira et al., 2007; Torrecillos et al., 2014). A computational model attempting to unify a broad range of findings on medial prefrontal cortex function (Alexander & Brown, 2011) proposes that this region is responsible for tracking discrepancies between expectations and outcomes, which are reflected in the FRN. Viewed in this way, the processing of execution and selection error may share a common neural network that signals a mismatch between the outcome and expectations in the service of behavioural adaptation (Cavanagh et al., 2012; Torrecillos et al., 2014).

To test whether outcome errors of action and selection can be dissociated in the medial frontal cortex, we recorded feedback-locked ERPs while participants engaged in a modified bandit task where choices were selected via rapid arm movements. Unrewarded trials were either framed as errors in choosing the wrong bandit (a selection error) or the result of an inaccurate movement (an execution error). Following a large body of evidence reporting that the FRN is sensitive to RPE (Sambrook & Goslin, 2015), we expected that unrewarded outcomes attributed to selection error would elicit an FRN response. If this medial frontal monitoring system also tracks general action-outcome discrepancies, then we should expect a deflection following errors of action execution too. However, should the recently proposed movement-dependent account of RL hold, the FRN response should be attenuated when errors can be ascribed to the motor system. We would expect P300 amplitude, a putative index of internal forward model revision (Krigolson & Holroyd, 2007a), to be largest for execution errors.

In addition to these predictions, we also examined the relationship between the FRN and behavioral modification. Specifically, we predicted that participants who exhibited a larger change in the FRN would be more likely to switch between the different options. Notably, we expected this brain-behavior relationship would hold for selection errors, but not for execution errors. Reasoning that action errors may instead be encoding information about the size of the execution error, with this feedback used to correct discrepancies between the planned and actual outcome, we explored the possibility that these signals may be correlated with the magnitude of error and subsequent change in motor response.

## Materials and Methods

### Participants

Using an effect size estimate derived from our previous work on the FRN (η2p = .167; Mushtaq et al., 2016), with a desired statistical power of 0.8 and alpha criterion set at 0.05, we set a minimum sample size of 28 participants. In total we tested thirty-two right-handed participants (EHI > 40; Oldfield, 1971). Two participants were excluded due to excessive EEG artifacts, and a technical error during data collection rendered one participant’s dataset unusable. All analyses were performed on the resulting sample of 29 participants (19 females, 10 males, µ age = 26.75 years, ±9.51 years).

Participants were told they would be remunerated based on their performance. However, due to the pseudo-veridical nature of outcomes (see Procedure), all received a fixed payment of £10.00. Participants signed an informed consent document, were fully debriefed, and the experiment was approved by the Ethics Committee in the School of Psychology at the University of Leeds, United Kingdom.

### Design and Procedure

We employed a novel three-armed bandit task (**Figure 1**) where the absence of reward on a given trial could be the product of a poorly executed action or an error in action selection (McDougle et al., 2019). Following EEG set-up, the participant was seated in a chair approximately 50 cm away from a 24” ASUS monitor (53.2 × 30 cm [2560 × 1600 pixels], 100 Hz refresh rate). The participant was instructed to make a choice by making a reaching movement, sliding their right arm across a graphics tablet (49.3 × 32.7 cm, Intuos 4XL; Wacom, Vancouver, WA) while holding a digitizing pen encased inside a customized air hockey paddle. The tablet was placed below the monitor on the table and between an opaque platform that occluded the hand.

**Figure 1.**
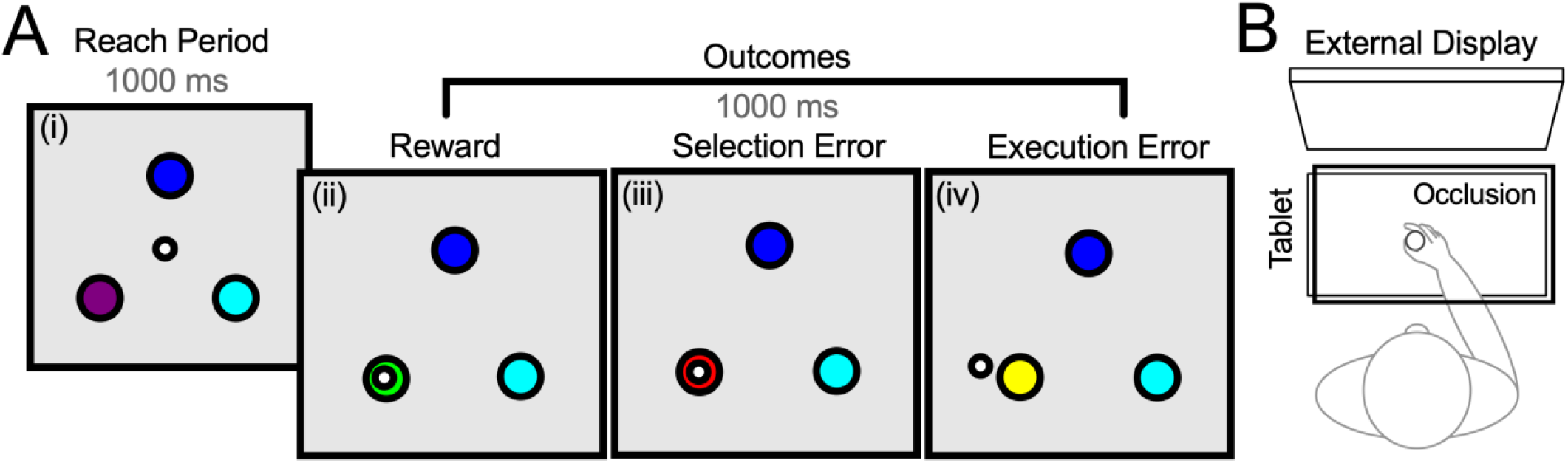
Experimental Task: (A) Participants moved a stylus on a tablet to make rapid shooting movements (i) through one of 3 bandits (large circles) at 90°, 210° and 330° degrees relative to the home position (small circle). Following a 1000 ms delay (not pictured), pseudo-veridical feedback (white cursor) was provided indicating if the outcome was a reward (ii), a selection error (iii) or an execution error (iv). (B) The hand was occluded throughout, and stimuli were presented on a monitor positioned in front of the participants at approximately eye level.

The experimental session comprised 400 trials, with opportunity for self-paced breaks. To initiate each trial, the participant made a reaching movement, sliding their right arm to position a white cursor (diameter of 0.5 cm) inside the home position, indicated by a solid white circle at the center of the screen. After maintaining this position for 400 ms, the start circle turned green and three bandits appeared on the screen (positioned at a radial distance 8 cm from the center at 90°, 210° and 330° degrees relative to the origin). The bandits were colored light blue, dark blue, or purple and the color-position mappings were maintained for the entire experiment (randomized across participants).

Following the appearance of the 3 bandits, participants had 2 seconds to initiate a reaching movement. If the reaction time (RT) was greater than 2 s, the trial was aborted and the message ‘‘Too Slow’’ appeared. After movement onset, participants had 1 s (Movement Time; MT) to complete a rapid straight-line “shooting” movement through one of the bandits. Upon movement initiation, the cursor indicating hand position disappeared and did not reappear until feedback presentation. If the movement was not completed within the required 1 s window, the trial was terminated and the error message “Too Slow” was displayed. If the movement was completed within the 1 s window, there were three possible outcomes: If the movement was accurate (hand passed through the bandit) the cursor was displayed within the spatial extent of the bandit. On these trials, there were two possible outcomes: (1) The bandit could turn green, indicating that a reward would be earned for the trial (reward outcome), or (2) the bandit would turn red, indicating that, while the movement was accurate, no reward would be given on that trial (selection error). If the movement missed the bandit, a cursor would appear indicating the position when the hand was at the radial distance of the bandits, and thus indicate if the execution error was clockwise or counterclockwise relative to the target. The bandit would turn yellow, further signaling an execution error. Participants were informed of the three possible outcomes prior to the start of the experiment and presented with demonstrations of the three outcomes.

Following McDougle et al. (McDougle et al., 2019), each bandit had its own fixed probabilities for the three trial outcomes. All bandits had a 40% reward outcome, and thus, the expected value for the three bandits were identical. However, the frequency of selection error and execution error trials varied. For one bandit, 50% of the trials resulted in execution errors and 10% resulted in selection errors. We refer to this as the “High Execution/Low Selection Error” bandit. A second bandit resulted in execution errors on 10% of trials and 50% resulted in selection errors (a “Low Execution/High Selection Error” bandit). A third, “Neutral” bandit produced an equal number (30%) of execution and selection errors.

To achieve these probabilities, outcomes were surreptitiously perturbed so that they aligned with predetermined feedback (a randomized sequence for each run) for the selected bandit. On trials in which the actual movement produced the desired outcome in terms of hitting or missing the bandit, the cursor was shown at its veridical position. However, if the participant’s movement missed the bandit, but the trial outcome was set as either a reward or selection error (i.e., outcomes requiring successful motor execution), the feedback showed the cursor landing inside the bandit, albeit near the side consistent with the actual hand position. Conversely, where a trial was set to be an execution error, but the stylus successfully intersected the bandit, the cursor was shifted just outside the bandit, with the side again consistent with the actual hand position (e.g., if the hit was slightly clockwise to the center of the bandit, the cursor appeared outside the spatial boundary of the bandit on the clockwise side). On trials in which feedback needed to be perturbed (i.e., deliver a false hit or false miss) to control the frequency of outcomes, the cursor position was shifted by randomly sampling from a normal distribution (± 6.24°, equivalent to .5 cm with an 8 cm reach) until a new cursor position was chosen that landed inside the bandit (for false hits) or outside the bandit (false misses).

We included three further constraints to minimize the likelihood that participants would recognize that the outcomes were not always directly reflective of their movements: (i) No online movement feedback was available; (ii) end-point feedback was presented 1 s after the stylus had passed the bandit location (this also helped reduce the impact of motor artefacts contaminating the ERP); and (iii) if the actual reaching angle was greater than 10° from the closest bandit on any trial (irrespective of the set outcome), no outcome was shown, the experiment software instructed participants to “Please Reach Closer to the Bandit.” Trials in which the movement was not completed within 1 s of the onset of the bandits or in which the reach angle was greater than 10° from the closest bandit were repeated, ensuring a full data set of 400 trials for each participant.

To increase motivation, participants were told that at the end of the experiment the software would randomly select five trials, and based on the outcomes from these trials, a cash bonus between £1-5 would be provided. As such, the goal was to accumulate as many reward trials as possible. In actuality, all participants received a fixed payment of £10 for taking part in the experiment.

Finally, given that it is possible that the execution error feedback could be interpreted in different ways (for example, participants may have assumed these errors were the result of faulty technical equipment), participants were invited to complete a brief optional post-experiment survey where they were asked to rate their agreement with the statement “I felt that that the miss (yellow) outcomes were the result of poor arm reaches” on a 7-point Likert scale, where a score of 7 indicated strongly agree and 1 indicated strongly disagree. From 21 respondents, a mean score of 5.57 (SD = 1.6), which was statistically significantly different to the mid-point (neither agree nor disagree) on the scale (t(20) = 4.41, p < .001), indicated general agreement with the intended experimental manipulation.

The experimental task was programmed using the Psychophysics Toolbox (Brainard, 1997; Kleiner et al., 2007) and lasted approximately 35 minutes, with an additional 25-30 minutes of technical set up for EEG data acquisition.

### Electrophysiological Data Recording and Preprocessing

EEG data were recorded continuously from 64 scalp locations at a sampling rate of 1024 Hz using a BioSemi Active-Two amplifier (BioSemi, Amsterdam). Four electrooculograms (EOG) – above and below the left eye, and at the outer canthi of each eye – were recorded to monitor eye movements. Two additional electrodes were placed on the left and right mastoids. The CMS and DRL active electrodes placed close to the Cz electrode of the international 10-20 system served as reference and ground electrodes, respectively. EEG pre-processing was performed using the EEGLAB (Delorme & Makeig, 2004) and Fieldtrip (Oostenveld et al., 2011) toolboxes, combined with in-house procedures running using Matlab (The MathWorks, Inc., Natick, Massachusetts).

All data were first re-referenced offline to the average of all channels, and downsampled from 1024 Hz to 256 Hz. The continuous time series data were filtered using a high-pass filter with a cut-off at 0.1 Hz (Kaiser windowed-sinc FIR filter, beta = 5.653, transition bandwidth = .2 Hz, order = 4638) and a low-pass filter with a cut-off at 30 Hz (Kaiser windowed-sinc FIR, beta = 5.653, transition bandwidth = 10 Hz, order = 126). A second filtering of the data was performed for subsequent independent component analysis using a high-pass filter cut-off at 1 Hz (Kaiser windowed-sinc FIR filter, beta = 5.653, transition bandwidth = 2 Hz, order = 4666). ICA typically attains better decompositions on data with a 1 Hz high-pass filter (Winkler et al., 2015). The data were segmented into epochs beginning 1s before and lasting 1s after the onset of feedback.

Infomax ICA, as implemented in the EEGLAB toolbox, was run on the 1 Hz high-pass-filter epoched data, and the resulting component weights were copied to the .1 Hz high-pass-filter epoched data. All subsequent steps were conducted on the .1 Hz high-pass-filtered data. Potentially artefactual components were selected automatically using SASICA (Chaumon et al., 2015), based on low autocorrelation, high channel specificity, and high correlation with the vertical and horizontal eye channels. The selections were visually inspected for verification purposes and adjusted when necessary. After removal of artefactual components, the Fully Automated Statistical Thresholding for EEG Artefact Rejection plugin for EEGLAB (Nolan et al., 2010) was used for general artefact rejection and interpolation of globally and locally artefact contaminated channels, supplemented by visual inspection for further periods of non-standard data, such as voltage jumps, blinks, and muscle noise.

Following artifact-removal, 93.5% of total trials were available for analysis. There was no difference in the percentage of trials removed across conditions (F (2, 56) = 2.09, p = .133). However, as a product of the experimental design, there was a difference in the total number of trials between the conditions (F (2, 56) = 85.2, p < .001), with more reward trials (µ = 150, ±9) available for analysis relative to execution error (µ = 114, ±12; t(28) = 12.21, p < .001) and selection error trials (µ = 110, ±11; t(28) = 13.89, p < .001). There was no difference in trial counts for the two types of errors (t(28) = .82, p = .693). To increase the reliability of our conclusions by addressing potential problems of distribution abnormalities and outliers, averaged waveforms were constructed for each individual by taking the bootstrapped (n = 100,000) means from the EEG time series epochs. The waveforms were baseline corrected using a 200 ms time window pre-feedback onset.

### ERP Quantification

Given that we had specific hypotheses, we focused our analysis on two locations. First, meta-analyses (Sambrook & Goslin, 2015; Walsh & Anderson, 2012) have shown the feedback-locked FRN effect to be maximal over the frontocentral region of the scalp. As such, we averaged activity across three frontocentral electrodes FC1, FCz, and FC2. Second, given that the P300 (specifically, the P3b sub-component) is commonly present in feedback-locked ERPs and typically maximal over parietal electrodes (Polich, 2007), we averaged over electrodes P1, Pz, and P2. Averaging across electrodes improves the signal-to-noise ratio of the ERP measures (Oken & Chiappa, 1986).

To test whether our results might be biased by the specific configurations of electrodes included in the averaged cluster and use of bootstrapped waveforms, we calculated the similarity between four different approaches to calculating the ERPs: (i) grand averaged activity from the raw waveforms in the clustered electrodes, (ii) grand averaged activity from the bootstrapped waveforms in the clustered electrodes, (iii) grand averaged activity from raw waveforms from a single electrode (FCz for frontocentral analysis and Pz for parietal); and (iv) grand averaged activity from bootstrapped means extracted from a single electrode. An intraclass correlation coefficient indicated a high level of agreement between all four approaches (Frontocentral ICC = .995, 95% CI 0.989-0.997; Parietal ICC: = .996, 95% CI 0.994-0.997). Clustered bootstrapped averaged ERP waveforms are reported here.

With growing evidence that most of the variation in the FRN is driven by a reward positivity, we decided to make use of difference waveforms for our analysis to detect differences irrespective of whether they were driven by positive or negative deflections in the ERP (Krigolson, 2018). A difference waveform procedure has the added benefit of more easily isolating the FRN from components that precede (P2) and follow (a large P3 component comprising a frontal P3a and parietal P3b), eliminating activity in common between two conditions (Kappenman & Luck, 2017). The majority of research on the FRN has typically computed “reward prediction error” (RPE) difference waveforms, derived by subtracting error/loss trials from reward trials (Sambrook & Goslin, 2015). Here, we created a “Selection Error” difference waveform by subtracting the average activity associated with Selection Error trials from the average activity related to all Reward trials, and an “Execution Error” difference waveform by subtracting the average activity associated with Execution Error trials from the average activity associated with Reward trials. Finally, we directly contrasted Execution and Selection Error ERPs by subtracting the Execution Error waveform from the Selection Error waveform to create an “Error Sensitivity” difference waveform. For statistical analysis, the parent waveform outcome trials were subjected to a one-way ANOVA and where main effects emerged, one-sample t tests were conducted to identify where these difference waveforms were significantly different to zero.

To reduce the number of false positives (Luck & Gaspelin, 2017), the ERP data were downsampled to 250 Hz and only activity between 150 and 500 ms (spanning the P2, FRN and P3 ERPs) was analysed. For each analysis, p values were corrected by applying a false discovery rate (FDR) control algorithm (Benjamini & Hochberg, 1995; Lage-Castellanos et al., 2010). The Benjamin-Hochberg correction approach was adopted as previous studies have shown it to reliably control the FDR when data are correlated, even when the number of comparisons are relatively small (Hemmelmann et al., 2005). This method is also ideally suited for the exploration of focally distributed effects (Groppe et al., 2011).

To aid the interpretation of the difference waveforms, we first visualised the grand averaged ERPs related to each outcome. For every statistically significant contrast, we present the mean amplitude from the cluster for each parent waveform. Differences between relevant conditions at each electrode site are also visualized through topographical maps to support interpretation of underlying components: Predicated on previous research (Walsh & Anderson, 2012), we anticipated that the FRN should show a frontocentral topography and, following an early frontocentral peak, there would be a subsequent posterior maximum corresponding to the P3b sub-component of the P300 (Holroyd & Krigolson, 2007).

### Brain-Behavior Relationships

A key question in this study is whether electrophysiological signatures of different types of outcomes correlate with the participants’ choice behavior (see San Martín, 2012 for a review). Based on a reinforcement learning account of the FRN (Holroyd and Coles, 2002), we would expect the amplitude of the FRN to scale with the degree of behavioral adjustment: large differences in the FRN should be more likely to lead to changes in choice behavior compared to small differences in the FRN. Here we can ask this question with respect to both selection and execution errors.

To examine brain-behavior correlations, we calculated a behavioral adjustment score, or “Switch Bias” rate, for each participant (operationalized as the ratio of the percentage of trials that the participant switched following an error to the percentage of switching following a reward). This served as an intuitive index of how much participants favored one outcome over another. Mean amplitudes from the statistically significant clusters of EEG activity were then correlated with these behavioral adjustment scores.

Rather than signaling a need to switch from one target to another, feedback from Execution Errors might be more readily used to modify a motor plan for future action. To quantify the magnitude of cursor error, we calculated the angular deviation of the cursor relative to the center of the selected target. Hand error was calculated as the position of the hand relative to the center of the selected target and was different to cursor error only on trials with perturbed outcomes. The degree of motor correction was examined on a subset of data where participants selected the same target on consecutive trials and quantified as the degree of angular change in hand position relative to cursor position on the previous outcome. Mean cursor error and motor correction scores were correlated with mean amplitudes from the previously identified statistically significant clusters of EEG activity.

### Statistical Analysis

For reporting purposes, time points are rounded to the nearest millisecond, amplitude (in microvolts; μV) to two decimal places and p values to three decimal places. The range for the scalp maps was time-interval specific and determined by the 1st and 99th percentile values across all electrodes. Spearman’s rho (r_s_) was used to examine correlations between amplitude and behavior. For correlations between behavior and neural activity, peak and mean amplitudes were extracted. Both are reported and the strongest correlations are visualized. Where appropriate, pairs of correlations were directly compared with Hittner, May, and Silver’s (2003) modification of Dunn and Clark’s (1969) approach, using a back-transformed average Fisher’s Z procedure as implemented in the R package Cocor v. 1.1‐3 (Diedenhofen & Musch, 2015). The statistical significance threshold was set at p < .05. Generalized eta squared (η_G_^2^) is used as a measure of effect size for repeated measures ANOVAs. This measure was selected over eta squared and partial eta squared because it provides comparability across between- and within-subjects designs (Bakeman, 2005; Olejnik & Algina, 2003); we considered η^2^_g_ = 0.02 to be small, η^2^_g_ = 0.13 medium and η^2^_g_ = 0.26 to be a large effect size. All statistical analyses were performed using R (R Core Team, 2015).

## Results

### Behavioral Responses

A one-way ANOVA revealed a significant difference in bandit preference (F [2, 56] = 8.27, p < .001, η^2^_g_ = .23), with participants exhibiting bias towards the High Execution/Low Selection Error bandit. Overall, this bandit was chosen on average on 39% (SE = 2%) of the trials, which was significantly greater than the Low Execution/High Selection error bandit (M = 29%; SE = 1%; t(28) = 4.03, p = .001) and Neutral bandit (M = 32%; SE = 2%; t(28) = 2.58, p = .046), with no difference for the latter two (t(28) = 1.07, p = .877). Consistent with previous work, when expected value is equal, the data show that participants prefer choices in which unrewarded trials are attributed to errors in movement execution rather than errors in action selection (Parvin et al., 2018; Green et al., 2010; Wu et al., 2009).

We then examined the effect of the different outcomes on the subsequent choice, asking how they influenced switching behavior (**Figure 2**A). Participants exhibited high switching rates overall (54%), but the rate differed according to outcome type (F [2, 56] = 10.23, p < .001, η^2^_g_ = .11). Switching was highest following selection errors (M = 66%; SE = 5%) and markedly lower following execution errors (M = 42%, SE = 5%; t(28) = 5.22, p < .001). This difference is consistent with the hypothesis that motor errors attenuate value updating, perhaps because participants believe they have more control to correct for execution errors (Parvin et al., 2018).

**Figure 2.**
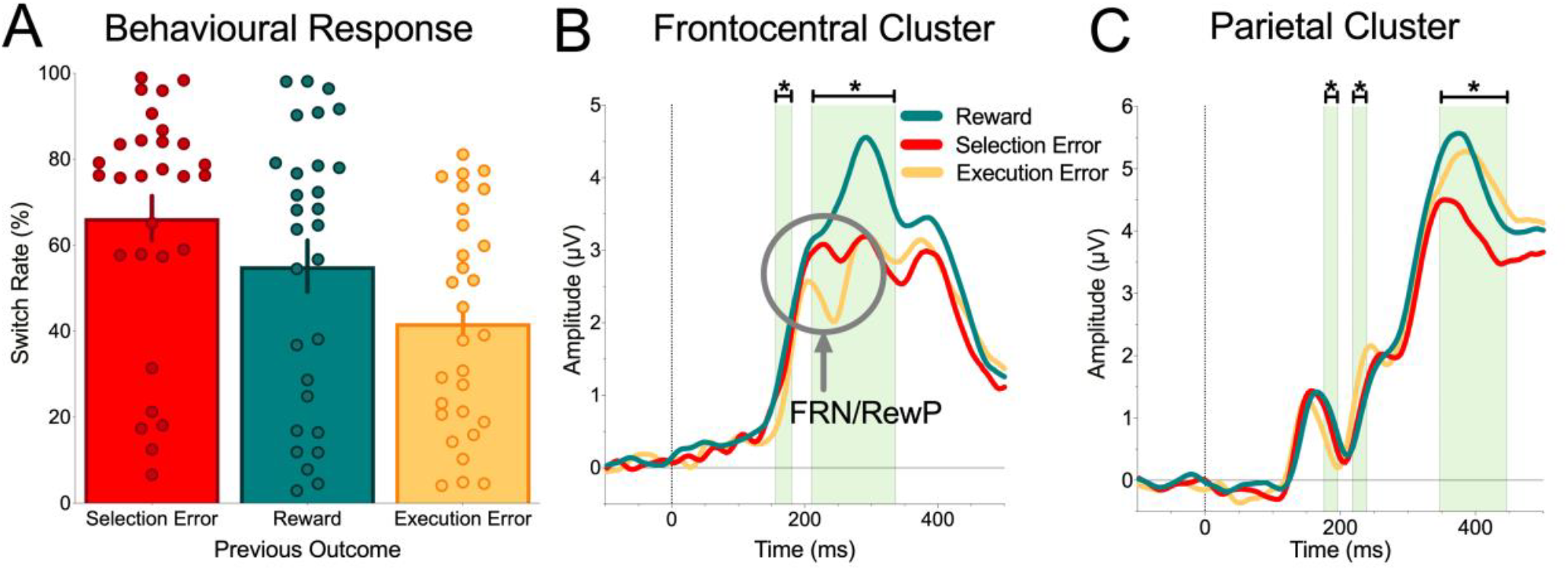
Behavioral Responses and ERP Grand Averages. (A) Switching rates following the three trial outcomes. Participants were more likely to repeat a choice (indexed by lower switch rates) following execution errors relative to selection error feedback. Error bars represent ±1 SEM. Feedback-locked ERPs for each outcome type, recorded from (B) frontocentral and (C) parietal electrode clusters. Zero on the abscissa indicates feedback onset. The green shaded regions indicate the significant clusters identified in the mass univariate analysis. Pairwise differences in these clusters are visualized in Figures 3-5 through the comparison of difference waveforms.

Interestingly, switch rates following rewarded trials fell between the other two outcome types (M = 55%, SE = 6%). There was no difference between switch rates following reward relative to selection errors (t(28) = 1.85, p = .227) or execution errors, although the latter approached significance (t(28) = 2.46, p = .062, following Bonferroni correction). The fact that many participants (18 of 29) were so prone to switching after a rewarded outcome and even more so (numerically) than after an execution error was unexpected. The high switching rates would suggest a bias towards exploratory behavior in this task-which might have been promoted by the relatively low rewards and/or the highly probabilistic nature of the outcomes (Cohen et al., 2007; Daw et al., 2006). Notably, there were very large individual differences in the treatment of the outcomes: Switch rates ranged from 3% to 98% following rewards, 7%-99% following selection errors and 4%-81% following execution errors.

### ERP Responses

Our primary aim was to examine whether selection and execution errors could be reliably distinguished in outcome-locked ERPs. To start, we ran an exploratory 3 (Bandit Type: High Execution/Low Selection Error vs. Low Execution/High Selection Error vs. Neutral) × 3 (Outcome: Reward vs. Selection Error vs. Execution Error) ANOVA at each time point for the frontocentral and parietal clusters. The main effect of Bandit Type was not significant (p’s ≥ .702) and there was no Bandit Type × Outcome interaction (p’s ≥ .671). Thus, we collapsed across the three bandits in our primary analyses of the three outcomes, allowing us to avoid increasing the family-wise error rate.

The grand averaged ERPs related to each outcome are shown in **Figure 2B and 2C**. F tests revealed two significant clusters in the frontocentral region between 156 -180 ms and 210-336 ms, and three clusters in the parietal region (176-196 ms; 218-239 ms; and 355-438 ms). Descriptively, the first cluster in the frontocentral region was driven by a delay in the onset of an initial P200-like signal following an execution error, and the second cluster incorporated FRN deflections following selection and execution errors, along with subsequent positive deflections, likely reflecting the P3a subcomponent of the P300 signal (Polich, 2007). The early two clusters in the parietal region reflect shifts in the latency and amplitude of the execution error ERP, with the third cluster driven by the attenuation of the P3b subcomponent of the P300 following selection errors.

**Figure 3A** depicts the Selection Error difference waveform, derived by subtracting the Selection Error waveform from Reward ERPs for the frontocentral cluster (shown in **Figure 2B**) and shows a statistically significant cluster of time points between 242-336 ms (one-sample t-tests of the difference wave against zero). An examination of the scalp topography of the first (242-289 ms) and second half of this window (289-336 ms) indicated a clear frontocentral maximum in the early phase, followed by a shift towards centroparietal maximum in the later part of the window (**Figure 3B**).

**Figure 3.**
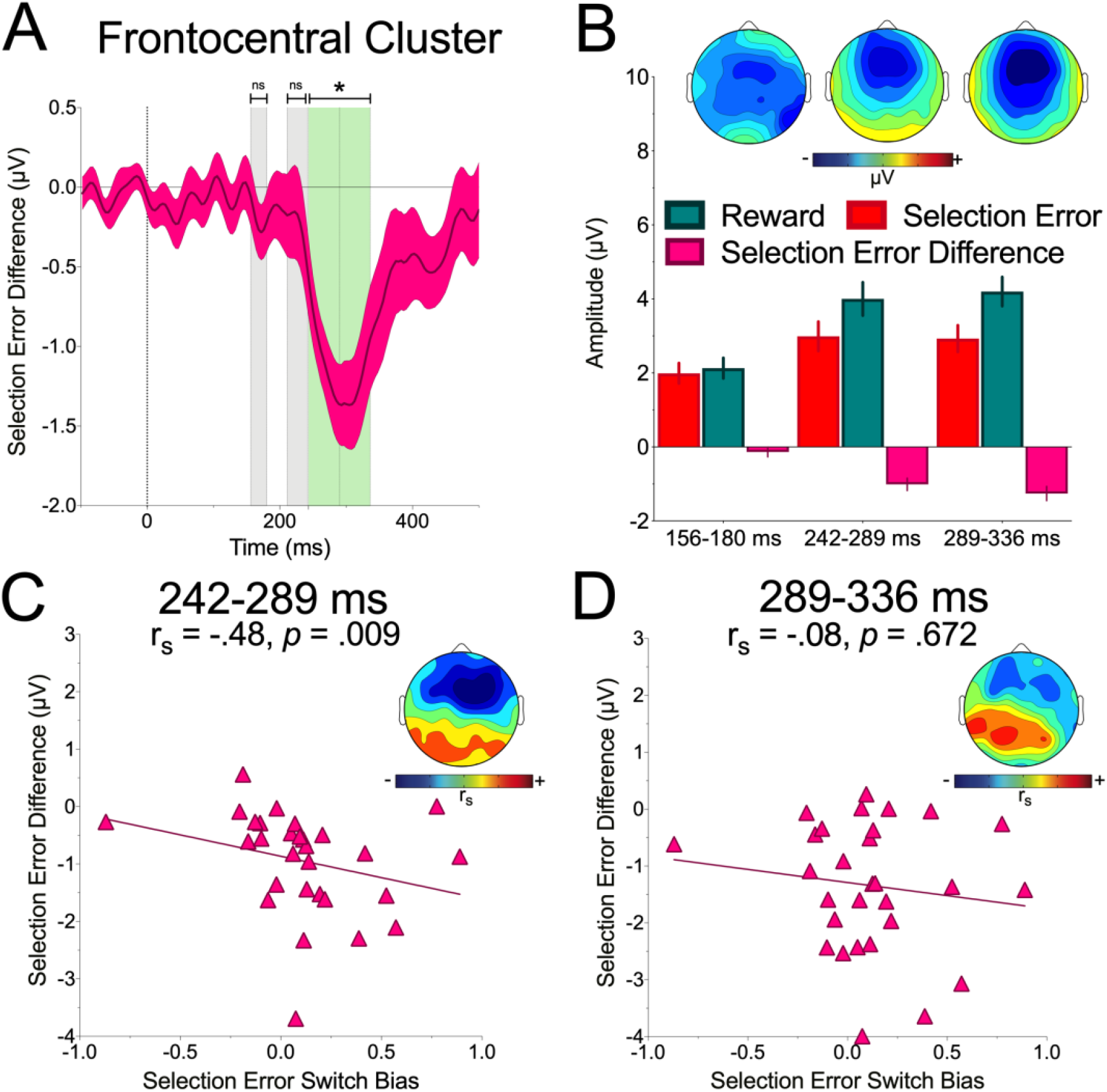
Selection Error in the Frontocentral Cluster: (A) The Selection Error waveform, defined as the difference in the ERPs on trials resulting in selection errors and rewards. The green shaded regions indicate significant clusters for this contrast and the grey shaded regions indicate where the clusters identified in the original time-series analysis did not reach statistical significance for this difference waveform. Zero on the abscissa indicates feedback onset. (B) Mean amplitudes for the early and late phases of the statistically significant clusters, with insets showing scalp maps of the distribution of differences across sites for each time interval. Selection Error difference waveform amplitude (shown on the ordinate, where negative values indicate more negative amplitude for selection errors relative to reward) correlated with an increase in the Switch Bias score (shown on the abscissa, where positive values indicate more switching following selection errors relative to reward) at a time interval corresponding to the FRN (C), but not the P3 (D). The insets show scalp maps of the distribution of amplitude differences across sites, revealing a frontocentral maxima for the FRN correlation.

In line with the reinforcement learning account of the FRN, there was a relationship between neural activity and behavior. Specifically, amplitude (mean: r_s_ = -.483, p = .009; peak : r_s_ = -0.36, p = .052; **Figure 3C**) from the early part of the cluster (capturing the FRN) negatively correlated with behavioral adjustment: The larger the difference waveform (i.e., greater negative deflection for selection errors relative to rewards), the greater the bias for the participant to switch to a different bandit following a selection error outcome relative to a reward outcome. We note that one participant had a switch rate score of -0.87, which was 2.97 standard deviations away from the mean. Re-running the analysis without this participant showed a weaker relationship, but the pattern remained statistically significant (mean: r_s_ = -.39, p = .042; peak: r_s_ = -.34, p = .074).

The topographical map (**Figure 3**C inset) demonstrates that this effect was localized to the frontocentral region. We found no evidence for such a relationship in the later, P3a, part of the time window (r_s_ = -.08, p = .672; **Figure 3**D). The mean FRN and P3a correlations were marginally different from one another (z = 1.96, p = .05), providing support that the FRN, but not the P3a, is a reliable correlate of behavior change.

### Execution Errors

To examine the electrophysiological correlates associated with unrewarded outcomes attributed to motor execution errors, we performed similar analyses, but now focus on the comparison between execution error trials and reward trials (the Execution Error difference waveform-the result of subtracting the Execution Error ERP from Reward ERPs in the frontocentral cluster shown in **Figure 2B**). This comparison revealed two statistically significant clusters- one ranging from 156-180 ms and a second between 207-325 ms (**Figure 4**A).

**Figure 4.**
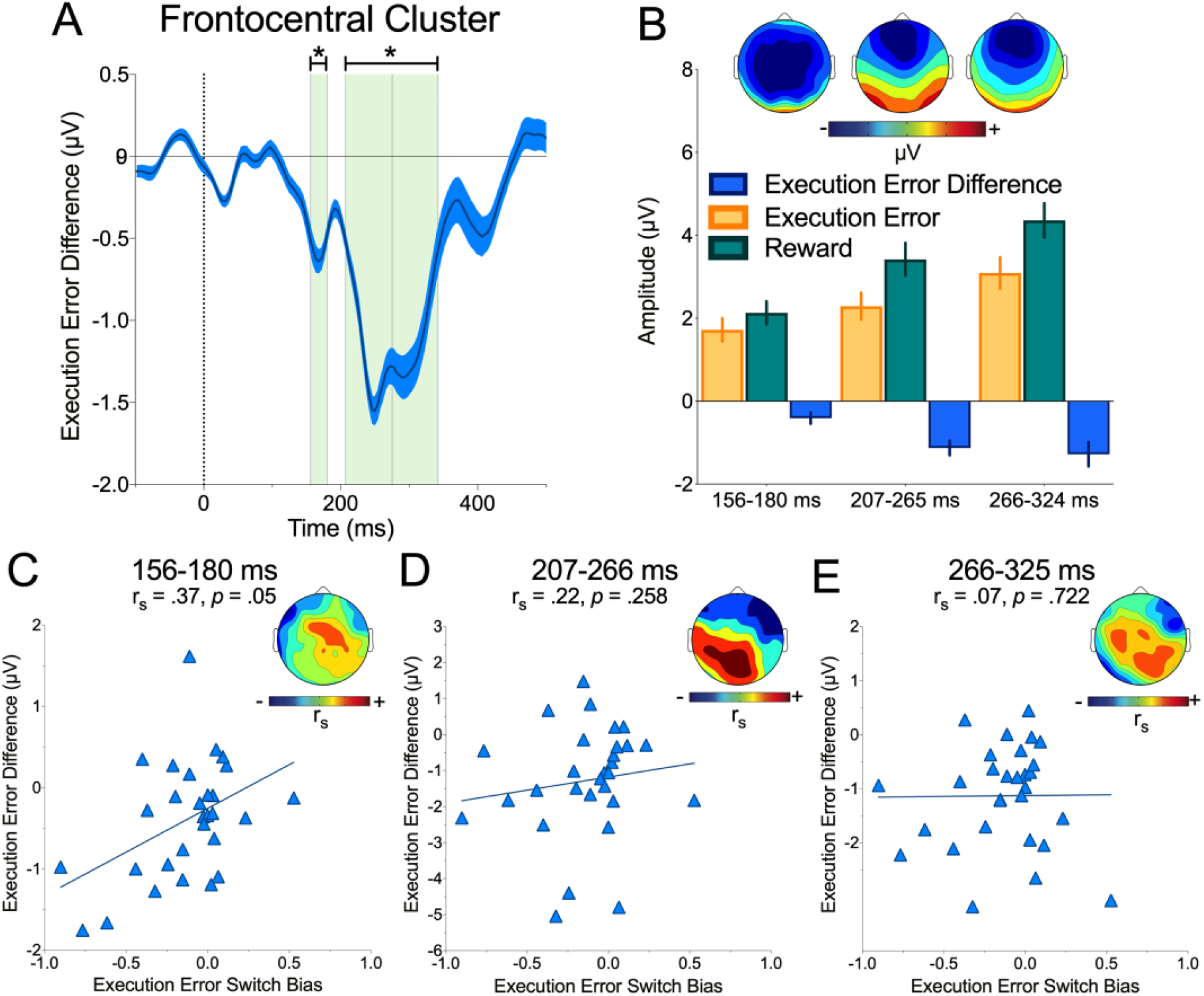
Execution Error in the Frontocentral Cluster: (A) The Execution Error difference waveform, defined as the difference amplitude for execution error and reward ERPs. The green shaded regions indicate clusters showing statistically significant differences. Zero on the abscissa indicates feedback onset. (B) Mean amplitudes for the early and late phases of the significant clusters. (C) The Execution Error difference waveform amplitude (shown on the ordinate, where positive values indicate larger amplitude for execution errors relative to reward) positively correlated with an increase in the Switch Bias score (shown on the abscissa, where positive values indicate more switching following execution errors relative to reward) in this early time window, but there were no correlations in the later time windows (D & E).

The first cluster showed an amplitude reduction in response to Execution Errors relative to reward trials. Similar to the Selection Error waveform result, we expected the second cluster would be contaminated by a P3a signal. Thus, we followed the same protocol, splitting this cluster into two equal intervals – (i) an early phase marked by the time interval 207-266 ms; and (ii) a later phase for activity between 266-325 ms. There was a clear frontocentral distribution for the early phase, and in the later time window, a shift towards centroparietal electrodes (**Figure 4**B).

We next examined the relationship between these three epochs (156-180 ms; 207-266 ms; 266-325 ms) and behavioral adjustment (**Figure 4**C-E). The peak amplitude difference in the earliest interval (156-180 ms) correlated positively (r_s_ = 0.37, p = .05) with switching rates following an execution error relative to reward. Following execution errors, smaller peaks in the 156-180 ms time window were associated with a lower tendency to switch. Note that this pattern is opposite to that observed between the amplitude of the FRN and behavioral adjustments following selection errors. The mean amplitude measure had a similar pattern of results, but was not significant (r_s_ = 0.35, p = .065). An examination of topography revealed this correlation to be maximal in the frontocentral cluster, suggesting that smaller amplitudes in response to execution errors early in the feedback processing stream are associated with a higher tolerance to this outcome.

In contrast to the results for Selection Errors, the FRN captured in the 207-266 ms time window did not correlate with behavioral adjustment (r_s_ = .07, p = .722). We tested, and confirmed, that this correlation was reliably different to the correlation observed for Selection Errors in the FRN time interval (z = 2.40, p = .016). There was no correlation between the Execution Error waveform in the P3a time window (266-325 ms) and behavioral adjustment (r_s_ =-.22, p = .258).

We conducted the same analysis for the Execution Error waveform in the parietal cluster of electrodes. Execution errors elicited smaller amplitude responses relative to rewards in an early time window (176-196 ms) but elicited larger amplitude responses at 218-239 ms post feedback. In the later time window, there was a positive correlation between amplitude and behavior (r_s_ = .47, p = .01) in the posterior region, suggesting a shift from frontocentral to parietal regions in the processes driving behavioral adjustment (Dhar & Pourtois, 2011; Overbeek et al., 2005). Interestingly, and unexpectedly, the amplitude of the P3b subcomponent of the P300 signal— proposed to reflect the revision of internal forward models in posterior parietal cortex (Krigolson & Holroyd, 2007a) showed no difference in the processing execution errors and rewards (see **Figure 2**C) and there was no relationship with behavioral adjustment (r_s_ = -0.01, p = .946).

### Error Sensitivity Difference Waveform

As described in the previous two sections, when using a common baseline (rewarded trials), we observed differences in both the ERP results and correlational analysis between unrewarded trials that were attributed to failures in movement execution or action selection. We performed a direct comparison between these two types of unrewarded outcomes by analyzing an Error Sensitivity difference waveform, subtracting the ERP for selection errors from the ERP for execution errors (see **Figure 2B** for the parent waveforms).

In the frontocentral cluster there was a significant difference in the range of the FRN (222-250 ms; **Figure 5** A, B). We had anticipated that the amplitude of the FRN would be attenuated following execution errors, assuming a lower response would be reflective of reduced value updating (McDougle et al., 2019). However, the observed effect was in the opposite direction: Execution errors elicited a larger FRN deflection, relative to selection errors.

**Figure 5.**
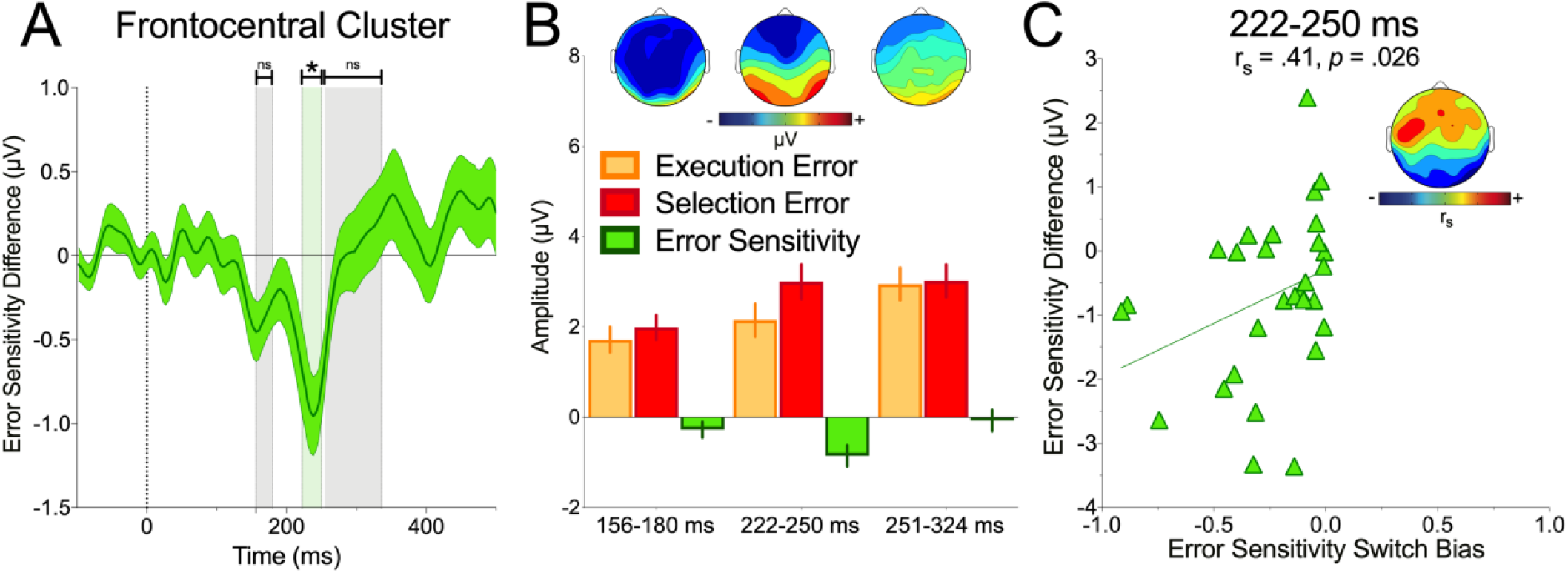
Error Processing Differences in the Frontocentral Cluster: (A) The Error Sensitivity difference waveform, calculated by subtracting ERPs for selection error from execution error ERPs. The green shaded region indicates the single cluster in which there was a significant difference for this contrast and the grey shaded regions indicate where the clusters identified in the original time-series analysis did not reach statistical significance in this comparison. Zero on the abscissa indicates feedback onset. (B) Mean amplitudes for the early and late clusters indicated by shaded regions in panel A. Inset scalp maps show topographical distribution for each cluster. (C) Peak amplitude difference in the FRN (shown on the ordinate, where negative values indicate a larger negative deflection for execution errors relative to selection error) correlated with a larger Switch Bias score (shown on the abscissa, where larger negative values indicate more switching following selection error relative to execution error). Note that no participants showed higher rates of switching following execution error relative to selection error. This correlation shows that as the similarity in the behavioral response to execution and selection error increased, amplitude differences in the processing of execution and selection error decreased.

We also examined whether the magnitude of this difference correlated with the “Switch Bias” rate. For this measure, the proportion of switches following execution errors was subtracted from the number of switches made following selection errors. Note that these values range from 0 to -0.91, due to the fact no participants produced more switches following execution errors relative to selection errors. Although the parent waveforms for this correlation are included in the previous analyses, the EEG activity in this analysis is specific to the range 220-250 ms, the window in which the error outcome ERPs differed significantly.

There was no relationship between mean amplitude in this window and Switch Bias (r_s_ = .23, p = .23). However, the peak negative amplitude revealed a positive correlation with Switch Bias (r_s_ = .41, p = .026; **Figure 5**C). Participants who had relatively similar switching rates to the two unrewarded outcomes had smaller FRN differences, while individuals with a large negative bias (i.e., less switching after execution errors) also exhibited larger FRN amplitudes for motor execution errors relative to selection errors. This correlation was maximal in frontocentral sites (**Figure 5**C inset).

Examining the parietal cluster revealed no differences in the earliest interval (176-196 ms).However, differences emerged in the 218-239 ms and 359-445 ms epochs, with larger positive amplitudes for execution errors relative to selection errors. The mean amplitude across each of these clusters (218-239 ms and 359-445 ms) was not correlated with the behavioral adjustment scores (r_s_ ≤ .179, p’s ≥ .352).

### Kinematic Analysis

To gain a deeper understanding of the relationship between brain activity and task performance, we examined correlations between task kinematics and the statistically significant periods identified in the time series analysis in the frontocentral and parietal difference waveforms. We reasoned that, in contrast to Selection Errors, where there was a relationship between FRN amplitude and choice selection, the Execution Error FRN may instead be encoding information about cursor position and subsequent movement correction.

In the first analysis, we examined whether there was a relationship between cursor error (the presented position of the cursor shown to participants at the end of the movement) magnitude and ERP activity. There were no reliable correlations between the mean activity of the statistically significant clusters in the difference waveforms and corresponding differences in cursor error magnitude (Execution Error: r_s_ ≤ 0.228, p’s ≥ 0.233; Selection Error: r ≤ 0.176, p’s ≥ .359; Error Sensitivity: r_s_ ≤ 0.152, p’s ≥ .429).

In the second analysis, we asked whether ERP amplitude on the current trial would correlate with the degree of motor correction on subsequent trials. Here, we restricted analysis to the subset of trials in which participants chose the same target consecutively. The amount of motor correction in response to feedback (computed as the mean absolute change in end-point veridical hand position relative to the cursor position on the previous trial), varied as a function of Feedback (F (2, 56) = 75.37, p <.001, η^2^_g_ = .66). As both outcomes indicated a successful movement, we expected, and found, no difference (t(28) = 0.47, p > .999) in the subsequent degree of correction for Selection Error (M = 3.73°, SE = 0.15°) and Reward (M = 3.64°, SE = 0.17°) trials. In contrast, Execution Error, signaling a need to change one’s motor response to hit the target (M = 6.53°, SE = 0.22°) had higher rates of correction relative to both Selection Error (t(28) = 8.95, p <.001) and Reward (t(28) = 8.95, p <.001) outcomes. Despite these behavioral differences, there were no correlations between mean activity of the statistically significant clusters in the difference waveforms and relative differences in the magnitude of subsequent motor corrections (Execution Error: r_s_ ≤ -0.239, p’s ≥ 0.211; Selection Error: r_s_ ≤ -0.328, p’s ≥ 0.083; Error Sensitivity: r_s_ ≤.152; p’s ≥ 0.429).

To ensure that we did not miss any potential sensitivity to task kinematics in other time ranges, we undertook an exploratory search of the full time series data by correlating cursor error and motor correction with mean amplitude from 150ms to 500ms.

We found no correlations between ERP difference waveforms and Cursor Error in the frontocentral (p’s ≥ .45) or parietal sites (p’s ≥ .75) following correction. We also note, with a degree of caution given the corrected p values were not significant, that there was one statistically significant pattern prior to correction- a positive correlation between the Error Sensitivity difference waveform and Cursor Error (r_s_ = .43, 406 ms). In correlating motor correction rates with ERP amplitude, we found no significant relationships in the frontocentral cluster (p’s ≥ .454). Here, we noted that the strongest relationship (r_s_ = .456) was a positive one between motor correction and the Error Sensitivity difference waveform at 164 ms – a pattern that was sustained across 156-174 ms. As participants made larger degrees of correction following Execution Errors relative to Selection Errors, they also exhibited greater amplitude. In the parietal cluster, we found no reliable patterns of activity following (p’s ≥ .97) or prior to correction (p’s ≥ .1).

### Perturbation Awareness

In a final set of explorations, we examined whether participants were sensitive to the feedback manipulation that had been applied to control the frequency of our three outcomes. In almost half the trials (M = 47.8%, SE = 0.01%) we delivered perturbed instead of veridical feedback (52.2%, SE= 0.01%). We had taken measures to minimize the likelihood of participants booming aware of these changes (e.g., no online movement feedback was provided, and end-point feedback was presented 1 s after the stylus had passed the bandit) and in a post-experiment survey, participants indicated that they believed execution error outcomes to be the result of poor reaches, suggesting no explicit awareness of the manipulation. Nevertheless, we did find differences in cursor error (**Figure 6**A), as revealed through a 3 (Outcome: Reward vs. Selection Error vs. Execution Error) × 2 (Veracity: Veridical vs. Perturbed) interaction (F (2, 56) = 27.4, p < .001, η^2^_g_ = .25). In all cases, cursor error was largest in the Veridical trials, but the effect was greatest for Reward (Veridical M = 1.68°, SE = 0.02°, Perturbed M = 0.98°, SE = 0.01°; t(28) = 26.83, p < .001) and Selection Error (Veridical M = 1.72°, SE = 0.02°, Perturbed M = 0.97°, SE = 0.02°; t(28) = 30.95, p < .001) outcomes, with differences of 0.7° and 0.75° respectively. For Execution Error, there was a visual difference of 0.27° (Veridical 5.99°, SE = 0.07°, Perturbed M = 5.72°, SE = 0.04°; t(28) = 3.5, p = .045).

**Figure 6.**
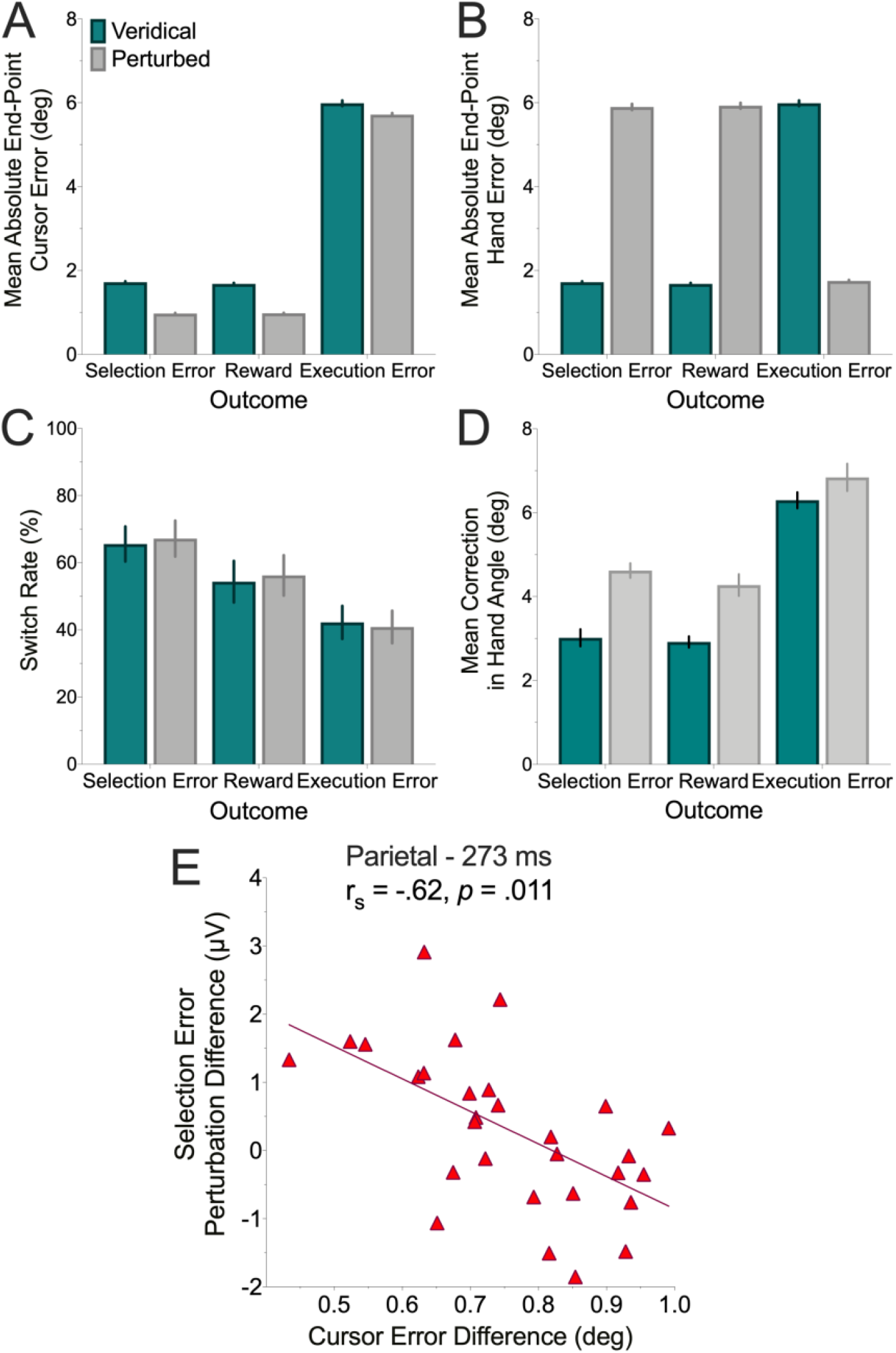
Feedback Perturbation and Awareness: (A) Cursor error was larger for veridical feedback relative to perturbed; (B) There was no difference in the magnitude of hand error for perturbed selection and reward error trials relative to veridical execution error trials and no difference between perturbed Execution Error trials compared to veridical Selection Error and Reward trials; (C) Despite smaller cursor error, participants made larger corrections in response to perturbed feedback, with the pattern most pronounced for false hits; (D) Perturbed feedback did not impact on the likelihood of switching bandits; (E) Amplitude differences between perturbed and veridical feedback in the Parietal cluster for Selection Errors at 273 ms (shown on the ordinate, where positive values indicate larger amplitude for veridical relative to perturbed outcomes) correlated with magnitude of the difference in cursor error for these outcomes (shown on the abscissa, where positive values indicate larger veridical cursor errors relative to perturbed).

In examining hand error (position of the hand relative to the center of the target), we found a Veracity × Outcome interaction (F (2, 56) = 4770.99, p <.001, η^2^_g_ = .981; **Figure 6**B). Veridical Execution Error trials (M = 5.99°, SE = 0.07°) were not statistically significantly different to perturbed Selection Error (M = 5.90°, SE = 0.07°; t (28) = 1.08, p = .886) and perturbed Reward trials (M = 5.93°, SE = 0.07°; t (28) = 1.09, p = .881). Similarly, there was no difference in hand error for perturbed Execution Error trials (M = 1.75°, SE = 0.02°) compared to veridical Selection Error (M = 1.72°, SE = 0.02°; t (28) = 0.998, p = .915) and veridical Reward trials (M = 1.68°, SE = 0.02°; t (28) = 2.41, p = .188).

Participants did not alter their behavioral strategy in response to feedback perturbations (Veracity: F(1, 28) = 0.899, p =.351, η^2^_g_ = < .01).; Veracity × Outcome: F(2, 56) = 1.42, p = .251, η^2^_g_ < .01; **Figure 6**C). However, a suggestion that they might have been implicitly sensitive to these differences is indicated by the degree of motor correction following veridical and perturbed feedback (**Figure 6**D). One participant had no stay trials following perturbed feedback in this subset of data and was excluded from this analysis. In the remaining participants, we observed an Outcome × Veracity interaction (F (2, 54) = 4.49, p = .016, η^2^_g_ = .04). There were no differences in the degree of motor correction following Execution Error (Veridical M= 6.3°, SE = 0.19°, Perturbed M = 6.84°, SE = 0.32°; t(27) = 2.07, p = .718), but greater corrections (Reward: Veridical M= 2.92°, SE = 0.13°, Perturbed M = 4.28°, SE = 0.26°; t(27) = 4.56, p <.001; Selection Error: Veridical M= 3.02°, SE = 0.20°, Perturbed M = 4.62°, SE = 0.17°; t(27) = 6.30, p <.001) followed false hits trials. These positively surprising outcomes (real reaches had missed the target on these trials, hence the perturbation) may have prompted overcompensation as participants sought to calibrate their movements to task feedback.

Given these differences, we explored the extent to which the ERP signal was sensitive to the veracity of the feedback. We re-ran the ERP time-series analysis, performing a 3 (Outcome: Reward vs. Selection Error vs. Execution Error) × 2 (Veracity: Veridical vs. Perturbed) at each time point for the frontocentral and parietal clusters. There were no statistically significant main effects of Veracity (F’s ≤ 6.99, p’s ≥ .397) and no Outcome × Veracity interactions (F’s ≤ 2.55, p’s ≥ .79) in the frontocentral cluster and similarly, no main effects (F’s ≤ 5.42, p’s ≥ .853) or Veracity × Outcome interactions (F’s ≤ 1.83, p’s ≥ .986) in the parietal cluster.

We then explored whether there were any differences in the relationship between ERP activity and kinematic adjustment as a function of Feedback Veracity. As perturbed feedback elicited larger corrective movements than veridical, we speculated that an ERP signal sensitive to positive surprise may scale in response to this behavior for Selection and Execution error trials. To explore this idea, a difference wave subtracting perturbed ERP amplitude from veridical was computed. The amplitude of this “Perturbation Difference” waveform was correlated with (i) the mean difference in cursor error for veridical and perturbed feedback per outcome; and (ii) the mean difference in degree of correction following veridical relative to perturbed feedback per outcome.

In analysing the relationship between the Perturbation Difference waveform and Cursor Error in the frontocentral cluster, we found no correlations that survived correction for multiple comparisons (p’s ≥ .616). However, in the parietal cluster, the Selection Error waveform strongly correlated with Perturbation Difference amplitude at 273 ms (r_s_ = -0.62, p = .011; **Figure 6**E), indicating a sensitivity to discrepancies between actual and presented hand position. Specifically, this correlation shows that for participants with larger veridical errors, perturbed feedback elicited larger positive amplitudes in a manner consistent with the P300 signaling surprise (Donchin, 1981; Nassar et al., 2019). The Error Sensitivity difference waveform showed a similar pattern but did not reach the significance threshold after correction (r_s_ = -.47 at 343 ms). The pattern for Execution Error was reversed, with the strongest correlation observed later (r_s_ = .45 at 492 ms)- with amplitude highest when both cursor error and amplitude were higher in the veridical condition relative to the perturbed condition. However, this too was not significant following correction.

In terms of the relationship between perturbation amplitude differences and the degree of motor correction, there were no significant effects in the frontocentral (p’s ≥ .120) or parietal clusters (p’s ≥ .82). With the same note of caution for non-significant correlations offered above, two patterns suggest a further dissociation in the processing of selection and execution error: In the time frame of the FRN, there was a relationship between frontocentral amplitude of the Perturbation Difference waveform and motor correction (r_s_ = -.542 at 289 ms). Here, greater corrective movements in response to perturbed feedback correlated with larger differences in the FRN; and (ii) later in the window, the Perturbation Difference waveform for Execution Errors positively correlated (r_s_ = .52 at 335 ms) with the degree of motor correction, indicating that larger cursor error corrections in response to perturbed feedback have correspondingly larger amplitudes for perturbed feedback in the time range of the P3a. Despite the finding that Selection Error, like Reward, resulted in adaptation following perturbed relative to veridical outcomes, no relationship was observed, with the strongest effect at 420 ms (r_s_ = -.299).

Finally, as an alternative to averaging over perturbed and veridical trials, we correlated the degree of perturbation on a single trial, computed as the difference between hand error and cursor error (which was zero on veridical trials, a positive value on trials where the cursor was shown to be closer to the target than the hand position and a negative value when the cursor position was shown to be further away from the target relative to hand position) with amplitude in the frontocentral and parietal clusters at each time point in the ERP per outcome for every participant. We did not find any general patterns to indicate a sensitivity to perturbation magnitude. In the frontocentral cluster, one participant showed a positive correlation between perturbation and the processing of Reward (between 152-172 ms and 254-289 ms), another showed a correlation for Execution Error trials (between 70-86 ms, 110-137 ms, 188-204 and 289-500ms) and two participants showed positive correlations for Selection Error. The first had a positive correlation between 453-457 ms and the second had a positive correlation in multiple clusters across the whole time series (between 4-11 ms, 31-90 ms, 117-188 ms, 258-277 ms, and 460 -477 ms). In the parietal cluster, no relationships emerged for Reward or Execution Error, with two participants showing positive correlations between the degree of perturbation and the processing of Selection Error: one between 340-356 ms and a second participant between 289-317 ms and 382-500 ms.

## Discussion

Adaptive behavior necessitates distinguishing between outcomes that fail to produce an expected reward due to either the selection of the wrong action plan or poor motor execution. Although the majority of decision-making research, in neuroscience as well as economics, have focused almost exclusively on the former, a few studies have shown that failed outcomes attributed to sensorimotor errors can markedly biases choice behavior (Green et al., 2010; McDougle et al., 2016, 2019). Here, we examined this issue by asking how an ERP signature of reinforcement learning, the Feedback-Related Negativity/Reward Positivity (FRN), varied in response to selection and motor errors. Predicated on the theory that the FRN is a scalp-related prediction error (Holroyd & Coles, 2002), we tested the hypothesis that errors attributed to failures in execution should lead to an attenuation in the FRN.

Consistent with our expectations, selection errors elicited a larger FRN relative to reward outcomes. Moreover, in line with a reinforcement learning account, the amplitude of the FRN following selection errors was negatively correlated with the probability that participants switched between the response options following feedback. Behaviorally, participants showed lower switch rates following execution errors, a pattern consistent with the hypothesis that the reinforcement learning system discounts these errors (McDougle et al., 2019). However, contrary to the prediction that FRN amplitude would be attenuated following execution errors, these errors actually produced the largest FRN. A striking difference between the ERPs in response to selection and execution error was that the amplitude of the FRN following selection errors was predictive of behavioral biases and learning, whereas this ERP response following execution errors did not correlate with these variables.

While almost all participants were more likely to switch after a selection error compared to an execution error, the differential response (i.e., difference in switch rates) to these two error outcomes varied considerably across participants. Moreover, this behavioral difference was correlated with the neural response to the two types of feedback: The more similarly participants treated the two outcomes at a behavioral level, the smaller the difference in FRN amplitude in response to these outcomes.

These findings could be reconciled by considering the top-down mechanisms that may modulate how execution errors are processed. Behavioral experiments have shown that a sense of agency related to the perceived ability to correct for motor errors biases choice behavior (Parvin et al., 2018). In the present experiment, the finding that participants persevered with a bandit following execution error but switched more often following selection errors also points towards differences in agency. Previous work on the FRN has shown that outcomes that can be controlled lead to a more negative FRN than those that cannot (Sidarus et al., 2017) and the FRN is attenuated in the absence of actively performed actions (Donkers et al., 2005; Donkers & van Boxtel, 2005). The finding that execution errors produced a larger FRN relative to selection error is consistent with the presumed greater sense of agency associated with this type of unrewarded outcome.

A recent fMRI experiment using a 3-arm bandit task similar to that employed here, revealed an attenuation of the signal associated with negative reward prediction error in the striatum following execution failures (McDougle et al., 2019). Our observation of a larger negative deflection for execution error trials in the FRN may appear contrary to these previously reported striatal results. However, the fMRI investigation did show increased ACC activity in response to execution errors compared to selection errors, suggesting that the former have their own neural signature. With regards to the EEG response, there have been a number of studies reporting FRN deflections in response to execution error (Anguera et al., 2009; Krigolson et al., 2008; Torrecillos et al., 2014). These studies, in line with the Prediction-Response Outcome model of medial frontal cortex function (Alexander & Brown, 2011), point to the existence of a general monitoring system that responds to violation of expectations. However, an important aspect of these tasks is that errors in movement execution typically resulted in high level goal errors (e.g., failure to reach or remain on target in a manual tracking task) and/or involved the introductions of perturbations during the movement phase (Krigolson et al., 2008). This makes it difficult to rule out the contribution of cognitive control and response inhibition processes-which are known to generate an N200 component that shares similar spatial and temporal characteristics to the FRN signal (Holroyd, 2004; Holroyd et al., 2008). A recent study separating reward and sensory prediction errors in a motor adaptation task showed that the FRN responds to the former, but not the latter (Palidis et al., 2019). The present findings, indicating qualitatively different relationships between the two medial frontal negativities with behavioral modification, add weight to the possibility that execution error processing may be distinct from dopamine-related reinforcement learning processes.

We also observed two distinct patterns of activity in time windows preceding and following the FRN that provide further support for the claim of differential processing of execution and selection error. First, smaller amplitude responses were observed following execution errors relative to rewards in frontocentral sites 156-180 ms post-feedback, and the amplitude of this component correlated with switch rates. Second, in parietal sites (218-239 ms), larger amplitude responses occurred following execution errors relative to reward and this difference was also correlated with switch rates. Importantly, in a reversal of the FRN pattern, magnitude differences in these early frontocentral and late parietal signals correlated with behavioral adjustment linked to execution errors. This pattern points towards the existence of distinct error monitoring systems operating at different levels of behavioural control (Yordanova et al., 2004).

Exploratory analysis on the relationship between ERP amplitude and task showed that the degree of motor correction following execution errors relative to selection errors correlated with amplitude differences in an early frontocentral cluster (156-174 ms). The time course of this cluster closely mirrored that of the earliest difference between execution error and reward – where amplitude differences correlated with switch rates. Given that we had no a priori expectations for such a result and that this specific result did not survive correction for multiple comparisons, interpretations must be treated with caution and require further robustly powered replication work to confirm. Should future work replicate this pattern it would add weight to the idea that the need to make a behavioural modification following an error in the motor system precedes the generation of the FRN.

A pertinent question of the present task and data is the extent to which participants were aware of the perturbations applied to the feedback to control outcome frequencies. Participants did not have access to online feedback and end-point cursor information was presented with a 1 second delay to minimize the likelihood of participants becoming aware of the perturbations. In a post-experiment survey, participants indicated that they had attributed execution errors to poor motor control. Consistent with this we found that during the task, perturbed feedback did not alter choice strategy, nor did it result in any significant differences in the ERP. However, participants did on average make larger corrective movements following perturbed feedback-this was despite these outcomes showing smaller cursor errors than veridical feedback. In exploratory analysis, we did not find any relationships between amplitude and perturbation magnitude at a trial level for the majority of the participants, but we did find a correlation between amplitude differences and cursor error when averaging across perturbed and veridical trials. This correlation manifested in the parietal cluster at 273 ms, which likely reflected the onset of the P300. Here, the positive amplitude of this signal reduced as the amount of veridical error increased. That the P300 shows a sensitivity to discrepancies between actual and presented hand position is consistent with the theory that the signal is generated through the active updating of an internal model of the environment (Donchin & Coles, 1988). The P300 is also notable for being a putative marker of conscious perception (Rutiku et al., 2015). If participants did indeed have access to this information during the task, it may be that these perturbations were not sufficiently large enough to signal a need to change strategy.

These findings also raise a broader question of whether the present results might be specific to outcomes that are framed as execution errors, or extend to any endogenous or exogenous event that results in an unrewarded trial in which the outcome does not provide information about the reward probability associated with the selected object (Green et al., 2010). For example, if an unexpected gust of wind blew a tennis lob out-of-bounds, would that be treated as an “execution error”? Or, if after pulling the lever on a slot machine, a power failure caused the game to terminate without a payoff, would this affect how the choice is judged? A future study could test endogenous execution errors (e.g., reaching error) and exogenous errors (e.g., the task screen goes blank randomly before an outcome is delivered) more explicitly than the perturbations applied here. If similar results are found in both settings, elements of the early activity observed in frontocentral sites may indicate the establishment of a sensory “state”, representing that the intended action plan was not properly implemented, irrespective of whether this mismatch was due to endogenous or exogenous factors, even before the prediction error is evaluated. This echoes the sequential ordering in models of temporal difference learning, where first the agent perceives its state, and then computes reward prediction errors relevant to that state (Sutton & Barto, 1998).

### Limitations and Future Directions

While we have hypothesized that execution errors impact choice behavior, either by attenuating the operation of reinforcement learning processes or via an enhanced sense of agency, it is also important to consider alternative hypotheses. In the behavioural data we observed a high base rate for switching between bandits. The highly probabilistic nature of the outcomes, coupled with the relatively low reward rate increased made the task of determining the optimal choice difficult (while each bandit different frequencies of execution and selection errors, they all had the same expected value). This may have biased participants towards an exploration strategy to reduce uncertainty by focusing on gathering more information about the reward likelihood of each bandit for later exploitation (Cohen et al., 2007; Daw et al., 2006). Viewed in this way, repetition of target selection following execution error might not be due to increased agency or RL discounting but may instead reflect a failure to acquire information on the reward probability of the chosen target on the previous trial and a drive to reduce uncertainty. Future work could disentangle these explanations by, for instance, assigning lower expected value to high execution/low selection error bandits and/or through the presentation of fictive outcomes for motor errors.

## Conclusion

We observed a robust FRN in response to both selection and execution errors, but only the former correlated with behavioral adjustment. In contrast, the amplitude of a positive deflection in the ERP, both prior and after the FRN, correlated with choice behavior following execution errors. These results indicate a need for a more nuanced interpretation of what the FRN represents, and how it may be shaped by contextual information. More generally, the results provide insight into how the brain discriminates between different classes of error to determine future action.

## Acknowledgements

The authors would like to thank Dr. Zeynep Uludag for her assistance with data collection and Dr. Assaf Breska for helpful discussions on an earlier draft of the manuscript. F.M and M.M-W were supported by Fellowships from the Alan Turing Institute and a Research Grant from the EPSRC (EP/R031193/1). J.A.T was supported by NxS084948 from the National Institutes of Health (USA). R.B.I and D.E.P were supported by NS092079 and NS116883 from the National Institutes of Health (USA). S.D.M was supported by F32MH119797 from the National Institutes of Health (USA).

